# Heterochronicity shapes brain network formation

**DOI:** 10.1101/2025.10.13.682136

**Authors:** Francesco Poli, Stuart Oldham, Alexa Mousley, Edward T. Bullmore, Petra E. Vértes, Duncan E. Astle

## Abstract

Brain network formation unfolds on a non-uniform developmental timetable, with different cortical regions generating connections at different developmental phases. Generative network models (GNMs) aim to uncover the principles underpinning the organisation of connectomes by creating synthetic networks according to simple computational rules. These models capture the connectome’s topology, operationalised here as the cumulative distributions of network metrics (e.g., modularity, small-worldness, rich-club structure). However, GNMs typically ignore the differential timing of connectivity formation, and by omitting this temporal programme they often misplace topological features in physical space. Here, we add a heterochronous growth term to GNMs and use a new model fitness function that weighs topology and topography equally. Topography refers to the spatial embedding of the network – the actual anatomical positions of the axonal tracts. With these advances, we can generate synthetic networks that more faithfully reproduce the spatial layout of diffusion-MRI connectomes from two independent adult cohorts. Compared with classical, temporally agnostic models, heterochronous simulations improve model fit, accurately locate cortical hubs and modules across individuals, and converge on a single caudal-to-rostral gradient of brain maturation. Overall, integrating heterochronicity makes GNMs more faithful to brain development, and allows for better identification of individual differences in network formation.

**Significance Statement:** A long-standing challenge in neuroscience is simulating brain networks that are anatomically realistic. By integrating a simple yet elegant principle – developmental timing, or heterochronicity – this study reproduces brain networks with an unprecedented spatial precision. We demonstrate that accounting for *when* different brain regions form connections is the key to accurately simulating *where* network features like hubs emerge. This shift from modelling static end-states to growth processes reveals a fundamental caudal-to-rostral maturation gradient. Our framework provides a biologically interpretable tool for identifying individual differences in brain development.

## Introduction

Understanding the structural organisation of the brain is crucial for explaining how distributed neuronal activity supports complex cognitive processes. By mapping the connectome, that is, the ensemble of white-matter tracts connecting distinct brain regions (1), we can chart the physical scaffolding that underpins information flow across neural circuits. In humans, this organisation is typically studied via non-invasive diffusion-weighted MRI, which reveals hallmark topological features such as a small-world architecture, hierarchical modularity, and rich-club organization (2–6). These features are thought to foster efficient communication among distributed brain areas, enabling the emergence of complex cognitive functions. During childhood, the ongoing refinement of large-scale networks accelerates the integration of distinct functional systems, laying the foundation for higher-order cognitive abilities such as language and executive control (7). By charting how network architecture emerges across time, we gain critical insights into the mechanisms that steer brain and mind maturation. In time, this may advance our understanding of how diversity in brain structure and function emerge, including in those cases where people encounter cognitive or clinical difficulties.

In the last twenty years, generative network models (GNMs) have become a crucial tool for understanding the architecture of brain networks (8–10). GNMs gain their explanatory power by compressing the connectome’s complexity into a concise, interpretable set of wiring rules. For instance, many connections can be explained by a “homophily” rule, whereby two brain areas are more likely to connect if they share similar connections to other areas (10–12). Concurrently, some penalties must be introduced to avoid generating connections that are too costly. For example, generating and maintaining long distance connections would require greater metabolic costs, and this should thus be avoided when possible (8, 13). In so doing, GNMs can reproduce many topological features of the connectome, such as its modular structure, rich-club architecture, and number of hub regions (2–6). GNMs provide a quantitative tool to understand which general rules any plausible wiring process must satisfy, improving also our ability to capture group-differences in disease (10, 14) and inter-individual differences (15). However, they are not intended to recreate the full biological mechanisms, or developmental timelines, that give rise to those networks.

While GNMs have successfully reproduced the topology of brain networks, they cannot capture their topographical properties (16). By topography, we refer to the spatial embedding of the connectome, including the exact physical locations of nodes and the spatial arrangement of modules and hubs. This is in contrast to the topology of the network which refers to the pattern of connections between nodes regardless of their spatial location. As a result, even though GNMs can correctly approximate the number of hubs, they often place these hubs in anatomically implausible positions. For example, models that prioritize wiring-cost minimization tend to pull hubs toward central positions, whereas in the empirical connectome they sit in peripheral cortical regions (17). This illustrates that there is an important conceptual difference between mirroring network topology and replicating the spatial or topographic reality of the brain.

We argue that existing GNMs fail to capture brain topography because their algorithms depart from the biological processes that sculpt neural wiring during development (15, 18, 19). In actual brains, the emergence of new connections during development is not random but follows spatially *heterochronous* patterns of neurodevelopment, which dictate the timing and manner in which neurons are born, migrate, and mature (20). Introducing heterochronicity into our models has the potential to capture these spatiotemporal constraints, thereby overcoming the limitations of existing GNMs. Ultimately, this approach could yield models that more faithfully approximate not only the topology of real brains, but also their topography (21, 22). Inspired by previous work (23), we mathematically define spatially ordered heterochronous developmental gradients. We hypothesise that this advancement will improve our ability to simulate brain structure. Moreover, this will in turn allow for the testing of hypotheses about the spatio-temporal principles that organise the brain.

One prominent theoretical question to be tested pertains to the ‘dual origins’ theory of brain organisation (24–29), which posits that the mammalian cerebral cortex has evolved along two principal gradients originating from the piriform cortex (paleocortex) and the medial wall, including the hippocampus (archicortex). These two foci of cortical expansion are expected to have a (partially) shared genetic program, but to give rise to anatomically distinct spatial patterns in terms of laminar thickness, neuronal density, and myelination (27). A second theory argues for different origin points, stressing how one neurogenesis gradient should start in the rostrolateral (frontal) region and proceed caudomedially (toward posterior sensory areas). The other origin point should be located in the thalamus, with primary sensory thalamic nuclei (lateral/medial geniculate, ventrobasal nuclei) maturing earliest and projecting to their respective posterior sensory areas sooner (25). By introducing additional parameters which vary the number and location of origin points, we can test for any existing model-based evidence in favour of either of these theories.

## Results

### Computational Advances

#### The energy function

We used GNMs to simulate brain network formation (Fig. 1A). These synthetic networks were then compared to real brain networks. Usually, the comparison is made using an energy function that considers how well the topological properties of the actual brain network are captured by the synthetic network. By energy, we refer to a single scalar “mismatch” score (lower is better) quantifying how unlike the synthetic and empirical networks are. In the classical formulation, energy is the maximum Kolmogorov-Smirnov (KS) distance between the cumulative distributions of degree *d*, betweenness centrality *b*, clustering coefficient *c*, and edge length *l*:

**Fig. 1.**
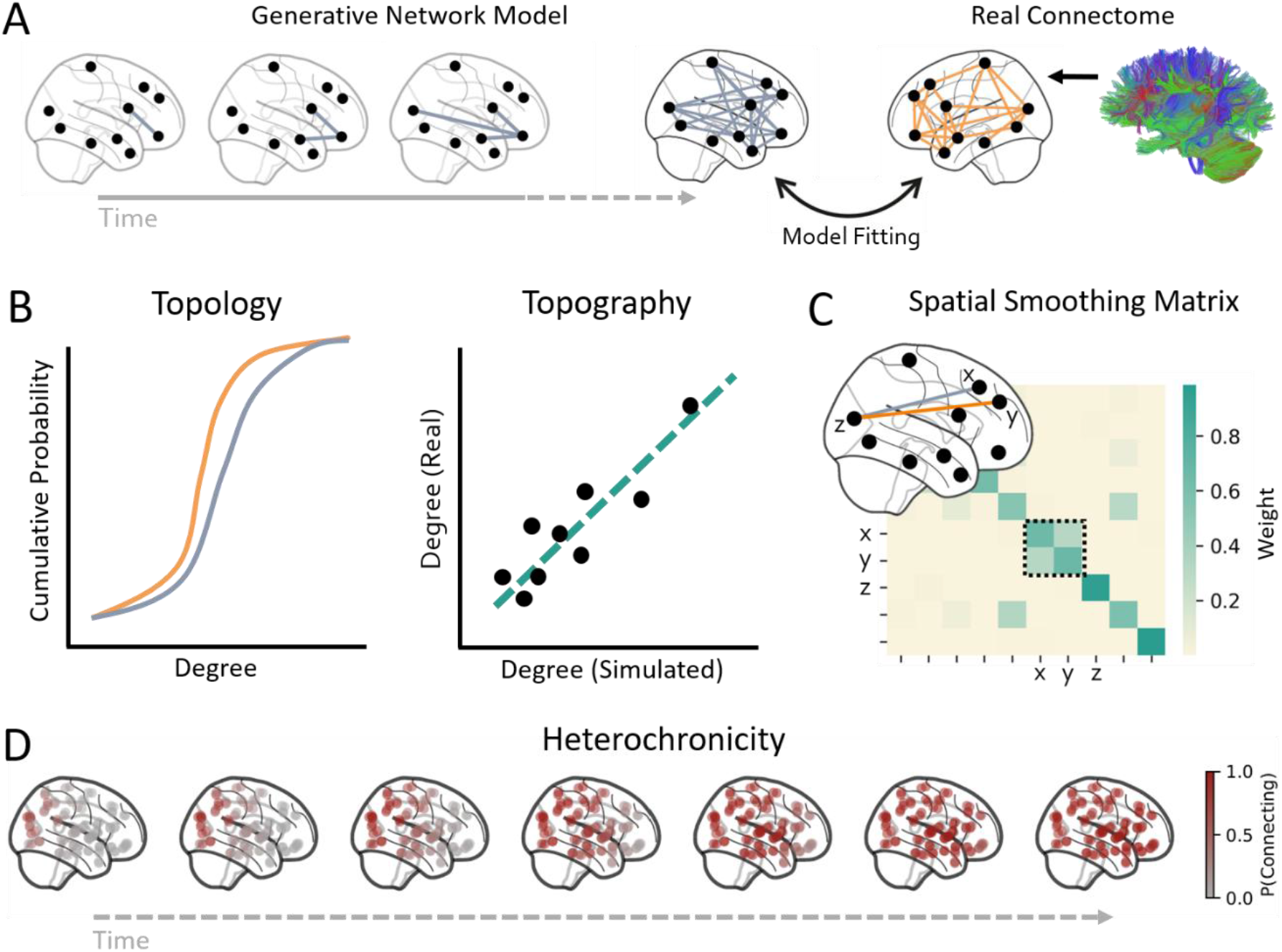
Generative network models simulate brain network formation. A. Starting from an empty graph, edges are generated one by one according to a cost–benefit rule that weighs Euclidean distance against a wiring value such as homophily. The process stops when the synthetic network reaches the density of the empirical connectome, reconstructed from diffusion MRI (streamline count), parcellated with the AAL atlas and binarized. Model parameters are optimised by minimising *energy*—the discrepancy between synthetic and empirical graphs. B. Conventional energy functions compare only topology, i.e. the overlap of cumulative distributions of node-level network metrics. We extend this criterion to include *topography*, i.e., whether corresponding nodes share similar network metric values. We assign equal weight to both components in the new energy function. C. The revised metric also grants partial credit to “near-miss” edges: if the model connects *z* to *x* instead of the empirical *z–y* link, the penalty is reduced when *x* and *y* are spatial neighbours. This is implemented by blurring every node’s metric with a distance-weighted (Gaussian) smoothing kernel so that neighbouring parcels share some of each other’s value. D. Example of heterochronous gradient starting from the primary visual cortex. The probability of a given area (each dot) of establishing connections with any other area is depicted in a grey-to-red scale.

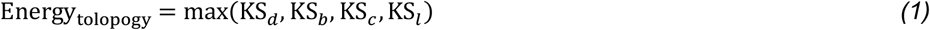

We expanded this energy score to consider topographical properties as well (Fig. 1B). That is, this addition considers not just the overall distribution of those topological properties (as in previous work), but also their spatial alignment with the empirical data. In other words, are hubs and peripheral nodes in the right place, and so on? More specifically, for each node we correlated its degree *d*, betweenness centrality *b*, and clustering coefficient *c* values in the synthetic versus real network, and took the lowest of those three correlations as the topographical cost:

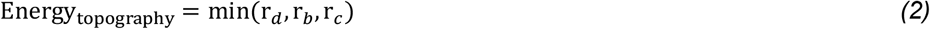

The resulting total energy was a sum of topological and topographical energies, each normalised such that they are weighted equally:

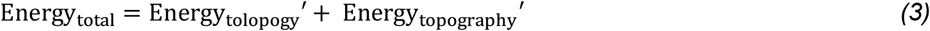

Since we aimed to capture the approximate (rather than the accurate) location of these network properties, we carried out a spatial smoothing before correlating the synthetic and real metrics (Fig. 1C). We used the new energy score to identify the GNM that produced the best model fit.

#### The generative model

Having defined how synthetic networks are scored, we now turn to the rule by which they are generated. The classic form of GNMs are composed of two terms, one for value (e.g., homophily) and one for cost (e.g., Euclidean distance) (10, 30). Here, we introduce a third term for heterochronicity:

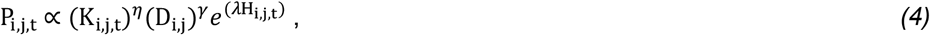

where P_i,j,t_ indicates the probability that a connection from node *i* and *j* is made at time *t; K, D* and *H* represent the matrices for homophily (i.e., matching index), Euclidean distance, and heterochronicity respectively, and γ, η, and λ represent three parameters regulating the extent to which the three matrices influence wiring probabilities. We refer to this new model as the heterochronous GNM, while the form without the 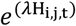 term we refer to as the classical GNM.

According to the heterochronicity matrix, H, different nodes start generating connections at different points in time. More specifically, we defined a Gaussian activation wave that started at an origin point (e.g., primary visual cortex in Fig. 1D) and gradually swept outward: Nodes closer to the origin increase their probability of connecting earlier, while those farther away start forming connections later. A formal definition of this heterochronous process is available in the Methods section under Computational Modelling.

### Heterochronous models produce better model fit than classical models

We fitted the GNMs using a grid search algorithm with γ, η, and λ as free parameters. Additionally, the origin point of the heterochronicity gradient was left free to vary across the medial axis of the brain, which naturally enforces symmetry. The models were fit to two datasets, the Cambridge Centre for Ageing and Neuroscience dataset (CamCAN) and the Human Connectome Project young adults dataset (HCPya) (see under *Datasets* in the *Methods* section) using the AAL atlas (31). All analyses reported below were additionally performed across an alternative parcellation (Schaefer100), an independent dataset (CALM), and an alternative tractography method (probabilistic; see Methods and Fig. S4, S5).

Next, a consensus network was derived for each dataset (32), thresholding the networks at 10% density (as in 14, 29). Given the stochastic nature of the generative process, each combination of parameters values generated 10 synthetic networks (i.e., 10 different runs with the same parameters values were performed) (33).

Results show that all parameters contributed to improve the total energy (Fig. 2A-C). More specifically, a sensitivity analysis (Fig. 2D; Fig. S1) showed that γ contributed the most to the decrease in total energy (CamCAN: β = -0.29, SE = 0.01, t = -19.99, p < 0.001; HCPya: β = -0.29, SE = 0.01, t = -20.82, p < 0.001), followed by λ (CamCAN: β = -0.14, SE = 0.01, t = -16.42, p < 0.001; HCPya: β = -0.14, SE = 0.01, t = -16.65, p < 0.001) and η (CamCAN: β = -0.06, SE = 0.01, t = -4.26, p < 0.001; HCPya: β = -0.06, SE = 0.01, t = -4.711, p < 0.001). Crucially, the energy of heterochronous models was significantly lower than the energy of classical models (i.e., when λ = 0) for both datasets (CamCAN: β = -0.08, SE = 0.02, t = -4.55, p < 0.001; HCPya: β = -0.06, SE = 0.01, t = -4.48, p < 0.001) indicating that introducing heterochronicity led to significantly more brain-like networks. More specifically, heterochronicity led to a shift in the trade-off between cost (i.e., distance) and value (i.e., homophily), with more negative parameter values for η and weaker positive parameters values for γ. This suggests that the heterochronous gradient actively promotes the formation of connections that are both homophilic and long-range. In turn, this alters the role of the other two model terms: because heterochronicity already drives up homophily, the homophily term η can be reduced without compromising topological fit. Conversely, because the gradient tends to generate more long-distance connections, the distance penalty γ must be increased to restrain the formation of metabolically costly links. In other words, the heterochronous mechanism shifts the balance of the model, requiring less homophily and more cost aversion to reproduce brain-like networks.

**Fig. 2.**
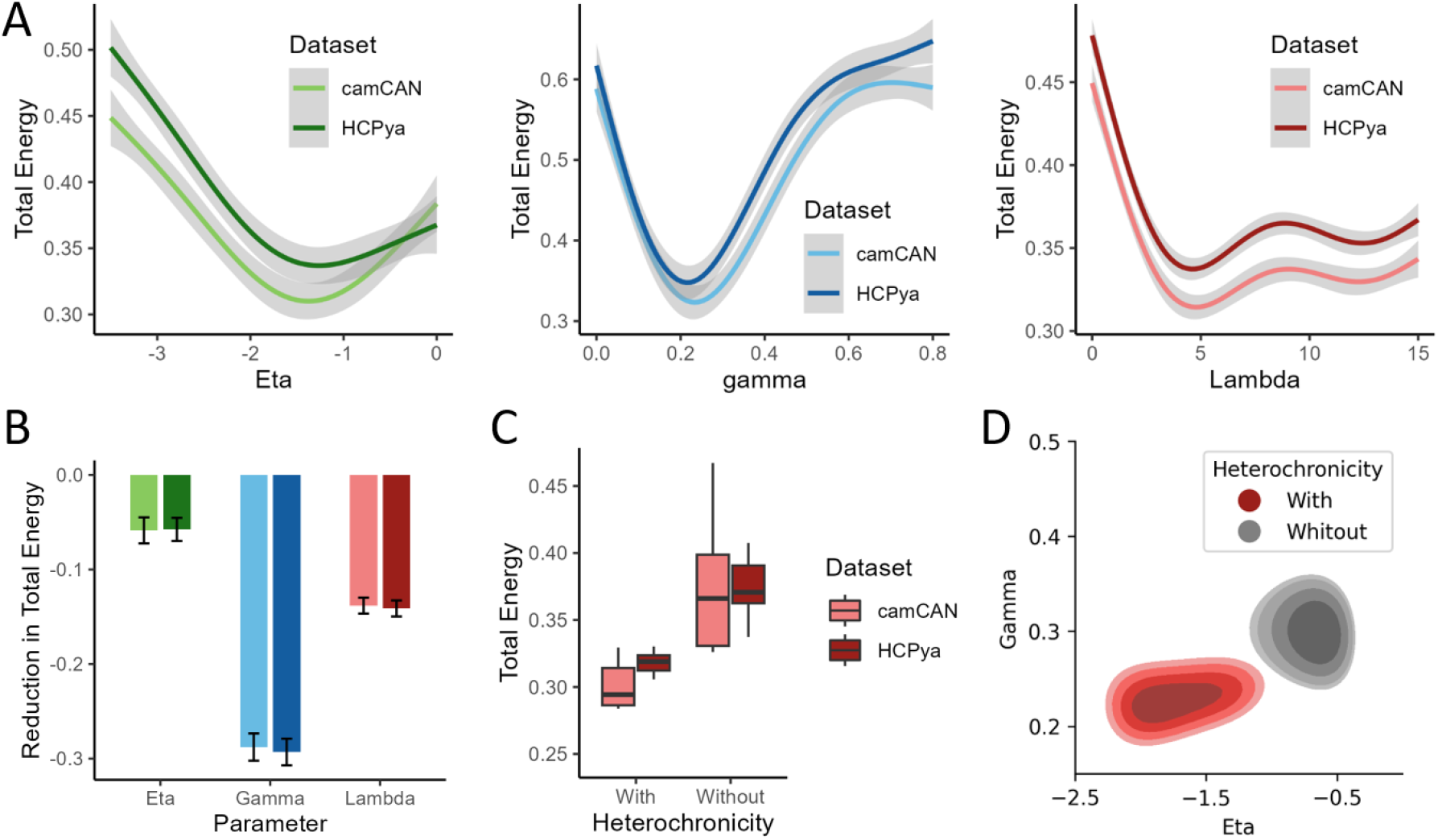
Homophily, heterochronicity, and distance penalty jointly contribute to generating plausible synthetic networks. A. Energy landscapes for each free parameter of the generative model, shown separately for the CamCAN (light) and HCPya (dark) datasets. For every panel, the remaining two parameters are fixed at the combination that yields the global energy minimum. Shaded ribbons denote 95 % confidence intervals across simulation runs. B. Global sensitivity analysis: standardised regression coefficients quantify how strongly each parameter contributes to lowering total energy (negative values indicate that larger parameter magnitudes reduce energy). C. Heterochronous models lead to significantly lower energy than classical models, indicating better model fit. D. Introducing heterochronicity leads to a shift in the trade-off between homophily (weakens) and distance penalty (strengthens).

Interestingly, per-participant fits revealed substantial individual variability in the parameter configurations producing the lowest energy (Fig. S2). Every individual was best fit by a heterochronous model (λ > 0; modal λ = 5), but with pronounced variability in heterochronicity strength and timing. Thus, while heterochronicity emerges as a robust feature of the best-fitting models, the strength and pace of the modelled gradient must be calibrated to each individual, raising the possibility that such variability captures genuine individual differences in developmental timing.

### Heterochronous models capture brain network topology and topography

Heterochronous models capture brain topology just as well as classical models, with low KS-statistics values for nodal degree (CamCAN: KS = 0.14, p = 0.30; HCPya: KS = 0.17, p = 0.12), betweenness centrality (CamCAN: KS = 0.14, p = 0.30; HCPya: KS = 0.17, p = 0.12), and clustering coefficient (CamCAN: KS = 0.17, p = 0.16; HCPya: KS = 0.12, p = 0.51). However, they perform *better* than classical models in terms of topography. More specifically, we found a positive correlation between the (smoothed) degree of the simulated and real network nodes (CamCAN: β = 0.45, SE = 0.01, t = 4.68, p < 0.001; HCPya: β = 0.35, SE = 0.01, t = 3.50, p < 0.001), their betweenness centrality (CamCAN: β = 0.40, SE = 0.01, t = 4.07, p < 0.001; HCPya: β = 0.25, SE = 0.01, t = 2.38, p = 0.019), and their clustering coefficient (CamCAN: β = 0.44, SE = 0.01, t = 4.66, p < 0.001; HCPya: β = 0.28, SE = 0.13, t = 2.18, p = 0.032). This is in stark contrast with classical models which lack heterochronicity, where no significant correlation between spatial maps of real and simulated nodes was found for clustering coefficient (β = -0.17, SE = 0.01, t = -1.63, p = 0.107) and betweenness centrality (β = -0.06, SE = 0.01, t = -0.53, p = 0.599), and this correlation was even significantly negative for nodal degree (β = -0.25, SE = 0.01, t = -2.46, p = 0.016). Overall, this indicates that introducing a heterochronicity gradient leads to a strong improvement in the model’s capacity to mimic brain network topography (Fig. 3).

**Fig. 3.**
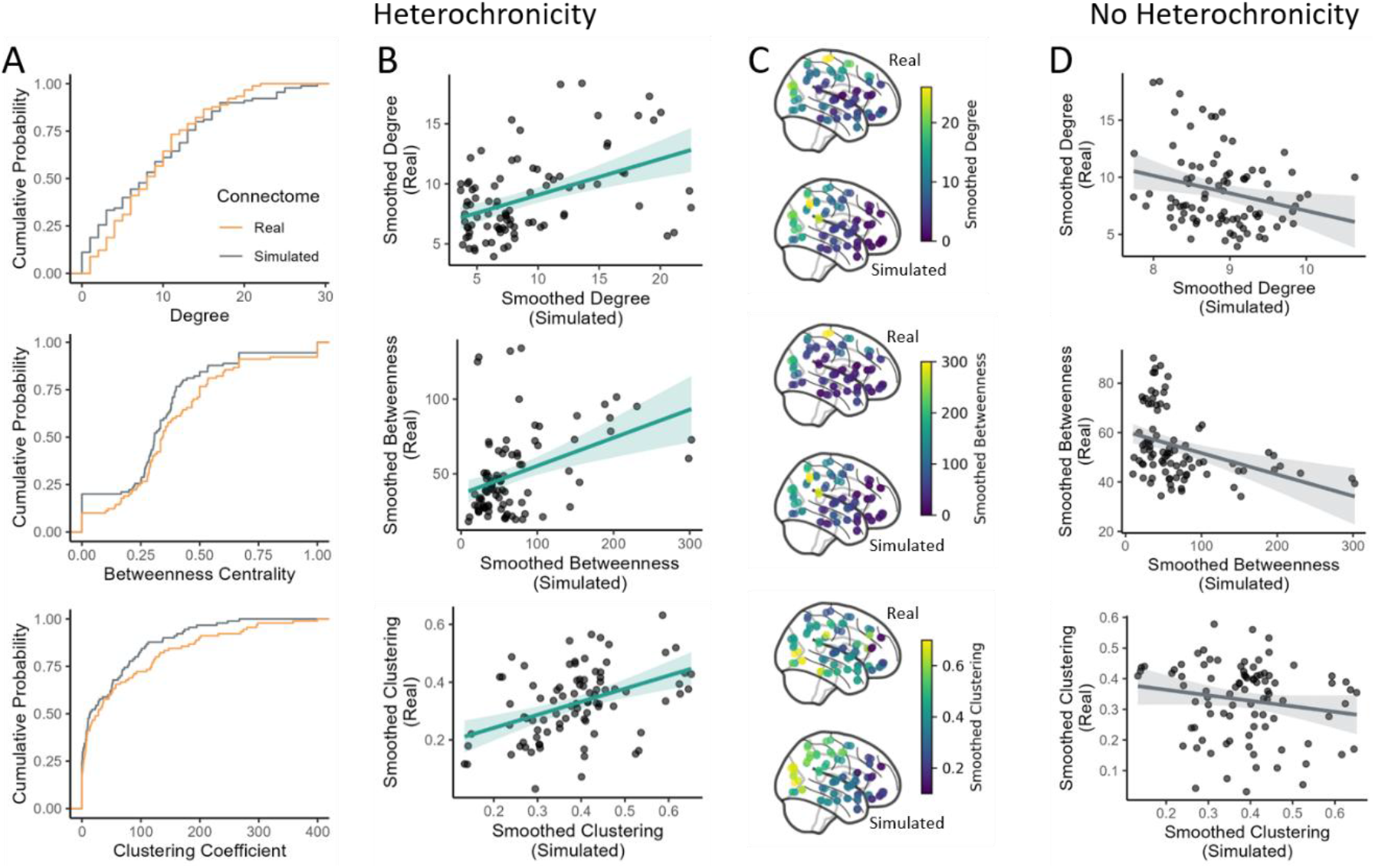
Heterochronous models capture real networks topography. A. Cumulative distributions of nodal degree, betweenness centrality, and clustering coefficient for the empirical connectome (orange) and its best-fitting heterochronous surrogate (grey). The two curves overlap closely for each metric, indicating that the model captures the global statistics (topological properties) of the real network. B. For the heterochronous model, vertex-wise values of the three smoothed metrics correlate with their empirical counterparts (solid regression lines, 95 % CI shading). C. Spatial maps of the smoothed metrics in the real brain (upper row of each pair) and the heterochronous surrogate (lower row) further illustrate their alignment. D. In contrast, the classical (non-heterochronous) model yields weak or negative correlations with the empirical data and fails to reproduce the observed spatial organization.

### Inferring the spatial origin of brain network formation

In our main analyses, the singular origin point of the heterochronicity gradient was free to vary across the medial surface of the cerebral hemisphere (Fig. 4A). Results from “sweeping” over the medial surface identified the more caudal part of the brain as the point of origin that reliably produced lower energy scores (Fig. 4B; Fig. S3) for both datasets. That is, when a single heterochronous wave is initiated at this point, the simulations are more brain-like. More specifically, we observe that more caudal origins led to lower energy (CamCAN: β = 0.04, SE = 0.002, t = 24.05, p < 0.001; HCPya: β = 0.04, SE = 0.002, t = 24.51, p < 0.001) and more dorsal origins also led to lower energy (CamCAN: β = -0.03, SE = 0.002, t = -16.66, p < 0.001; HCPya: β = -0.04, SE = 0.002, t = -22.26, p < 0.001).

**Fig. 4.**
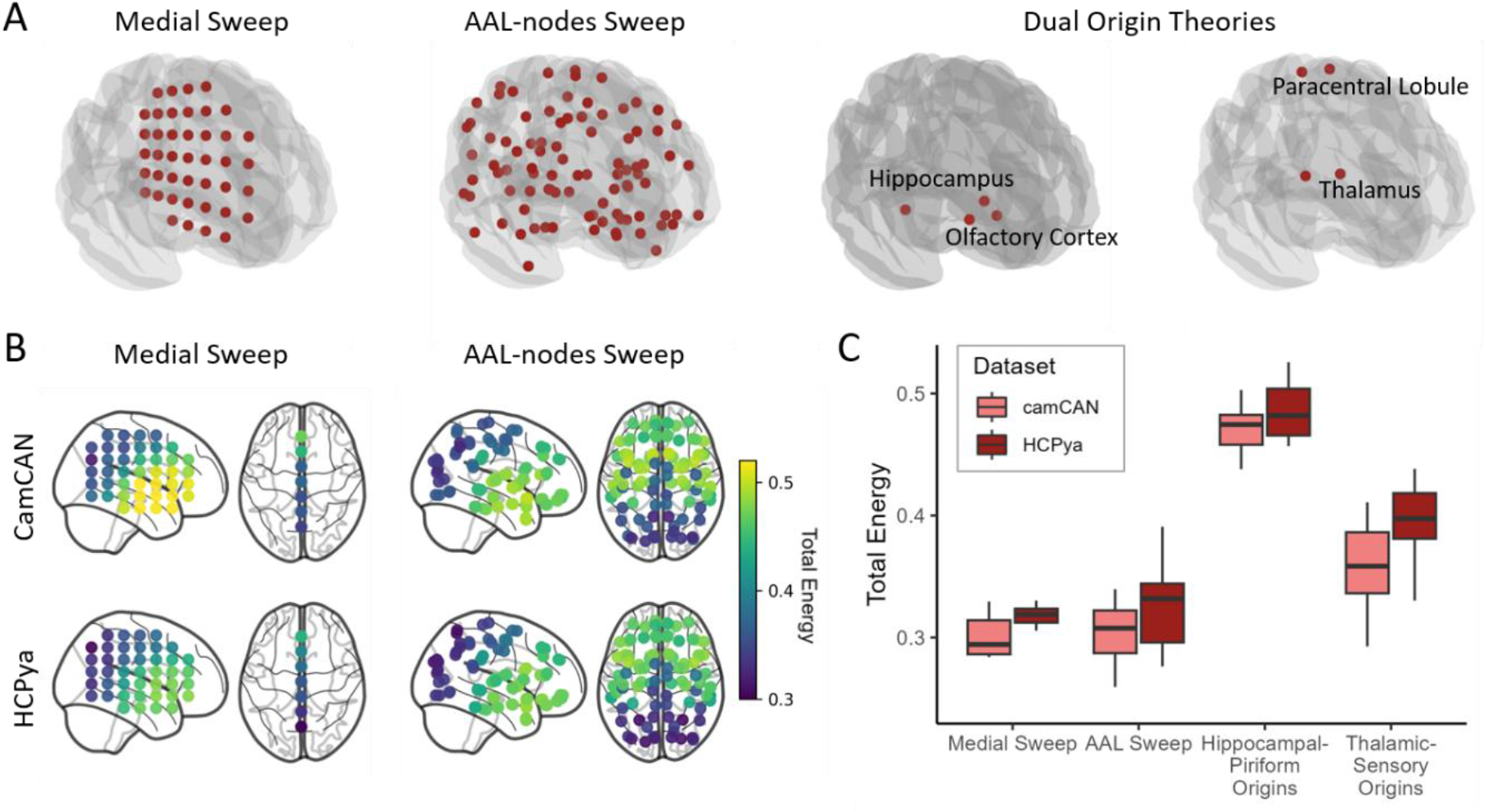
Single-origin heterochronous growth seeded from dorsocaudal cortex best reproduces real networks. A. Seed configurations used to initialise heterochronous growth models. Red spheres indicate (i) a systematic “medial sweep”, (ii) a sweep across anatomical AAL nodes, and (iii) two dual-origin hypotheses (hippocampal-piriform and thalamic-prefrontal pairs). B. Vertex-wise total energy for the medial and AAL sweeps in both datasets. Seeds placed farther dorsally and caudally yield lower energy, implying a closer match to the empirical network’s topology and topography. C. In both datasets, single-origin models outperform dual-origin models; Lower energy indicates a more accurate synthetic network.

To further probe potential origins of brain network formation, we run additional models manipulating the origin points. First, we performed a sweep over the *x-y-z* coordinates of all AAL nodes. To maintain hemispheric symmetry, the point-of-origin was never located in a single hemisphere, but always bilaterally. Results from the sweep over the AAL coordinates replicated the initial results, identifying the caudal part of the brain as the origin point that reliably produced lower energy scores for both datasets (Fig. 4B). Thus, our model indicates the heterochronicity gradient generally moves from caudal to rostral direction.

Second, we tested specific dual origin theories. One theory focuses on the dual origin of the cortex, pointing at hippocampal and olfactory cortices as origin points (27); the other focuses on the entire brain, pointing at the thalamus and sensorimotor cortices as possible origin points (Finlay & Uchiyama, 2020; Fig. 4A). When looking at the energy produced by the dual origin models, we saw significantly higher values for both the hippocampal-piriform origin (CamCAN: β = 0.14, SE = 0.02, t = 9.06, p < 0.001; HCPya: β = 0.13, SE = 0.01, t = 9.68, p < 0.001) and the thalamic-prefrontal origin (CamCAN: β = 0.13, SE = 0.01, t = 8.67, p < 0.001; HCPya: β = 0.14, SE = 0.01, t = 10.70, p < 0.001), indicating poorer model fit. Hence, at least with the current data type and pre-processing steps (i.e., streamline counts, parcellated in AAL), we do not find evidence in favour of a dual origin of the brain.

### Advances and systematic limitations in charting long-range connections

Models incorporating heterochronicity create networks that are more topographically similar to real brains, relative to classical models, but are they doing any better in the number and placement of critical long-range connections? This was a noteworthy failure of classical models (16). To test whether heterochronous models led to any improvements in this regard, we checked differences in the distribution of the lengths of the connections between the real, the standard, and the heterochronous connectomes (Fig. 5A-B). By comparing KS-statistics across all runs of the models, we find that the heterochronous connectomes are more similar to the real connectome than the classical ones for CamCAN (β = 0.15, SE = 0.006, t = 24.65, p < 0.001), although this difference was not significant for HCPya (β = 0.007, SE = 0.007, t = 1.00, p = 0.324). This improvement was specifically due to the right tail of the distribution, which concerns long-range connections (see Fig. 5A). Hence, the heterochronous model better simulates the correct proportion of long-distance connections, relative to standard GNMs.

**Fig. 5.**
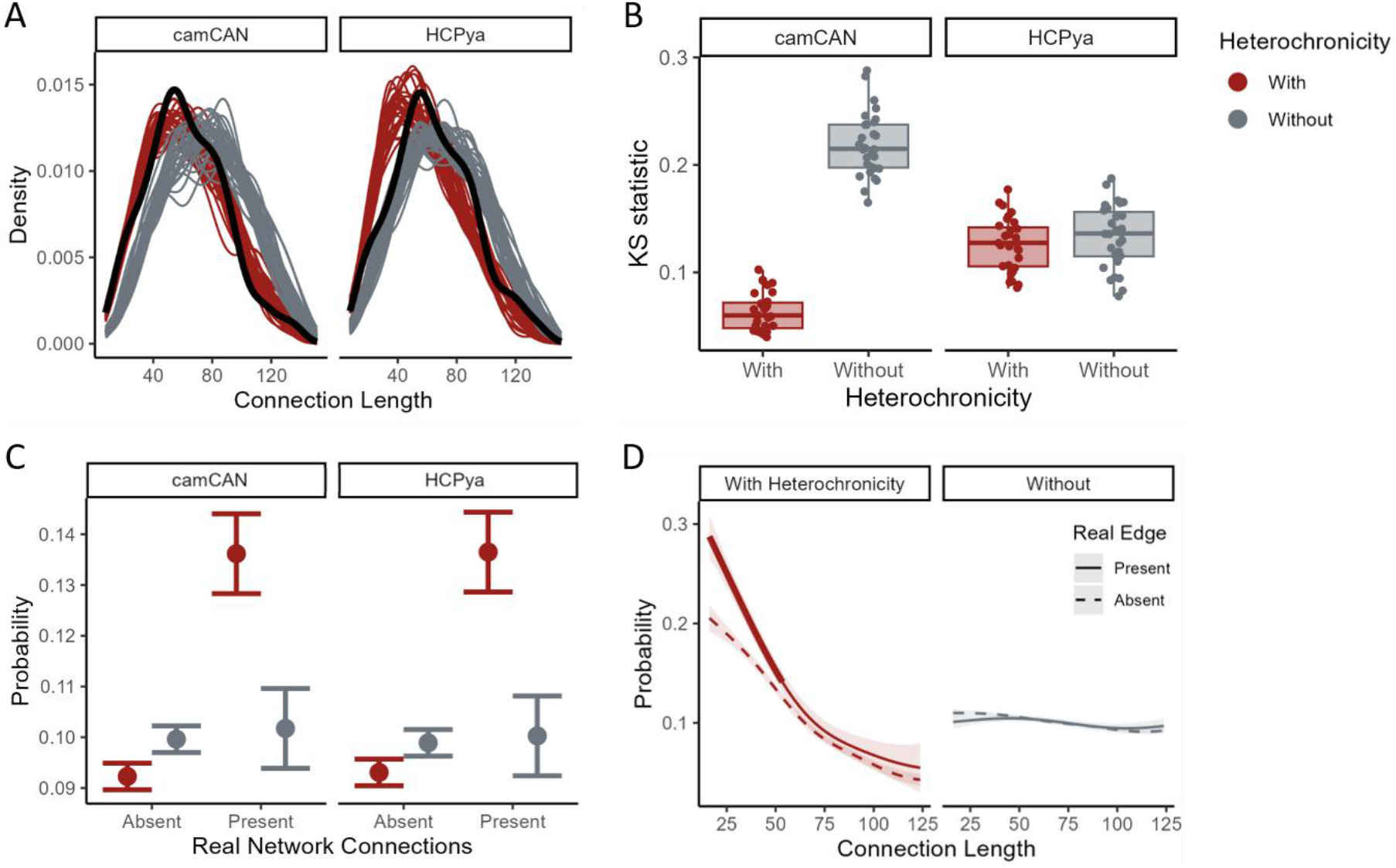
Heterochronous models place correctly only short-range connections. A. The distributions of connection lengths generated by the heterochronous model (red) and the classical model (grey) across individual simulation runs, plotted against the empirical consensus network (black). B. Two-sample Kolmogorov-Smirnov (KS) statistics comparing each synthetic network with the empirical distribution. Lower values indicate a closer match. The heterochronous model fits the CamCAN data significantly better than the classical model, whereas no clear difference is observed for the HCP-YA dataset. C. Mean probability of drawing edges that are present (right) or absent (left) in the empirical network. In both datasets, heterochronous models outperform classical models at identifying existing connections and rejecting non-existing ones. Error bars show 95 % confidence intervals. D. Probability of drawing edges that are present or absent in the real connectome as a function of connection length. The heterochronous model is accurate for short-range connections (thick red line) but falls to chance for long-range connections, whereas the classical model remains at chance across all distances.

Heterochronous models do not simply generate a more faithful distribution of the real connections but are also more likely to place the connections in the right place. Given that, compared to classical models, heterochronous models are more likely to place connections where they actually exist, and less likely to place them where they do not exist (Fig. 5C). However, they perform above chance only for short- to mid-range connections. To show this, we compared the model’s probability of placing a connection where one truly exists against where it does not as a function of the connection length (Fig. 5D). Heterochronous models assigned a higher probability to connections that truly existed (gap = 0.021, 95% CI [0.013, 0.028], p < 0.001), but this advantage was significant only for short- to mid-range connections (∼16–53 mm); classical models showed no difference at any length (gap = −0.001, p = 0.42). This systematic limitation will be addressed in the discussion.

## Discussion

Incorporating a spatially heterochronous developmental gradient into generative network models markedly enhances our ability to recreate not only the topology but also the topography of human brain networks. By constraining nodes to make connections following a probabilistic spatiotemporal gradient, the model produces synthetic connectomes whose spatial distributions of graph metrics align far more closely with empirical diffusion-MRI data than classical, temporally agnostic models. This improvement underscores the importance of incorporating temporal aspects of development within computational models of the adult connectome. Likewise, computational models offer a way of empirically formalising and testing developmental theory.

The parameter search consistently pointed to a caudo-rostral wave of connectional maturation: a single point of origin located in posterior medial cortex yielded the lowest energy across datasets. This caudal origin point gave occipital and parietal regions more time to connect with other areas, which in turn led them to become high degree hubs in the simulated adult network. This is consistent with evidence indicating that occipital and parietal regions show more connections in human cortical networks (34, 35); and with previous work showing earlier onset and therefore more prolonged myelination of occipital compared to prefrontal areas (36).

At first glance, the result conflicts with accounts that posit a dual brain or cortical origin (e.g., hippocampo-piriform or thalamic-sensorimotor). However, it is important to delineate the purposes of those theories, that is, to explain neurogenesis and laminar differentiation. In contrast, our model is a computational formalisation of the later-emerging pattern of white-matter connectivity. It remains entirely plausible that neurons are born in multiple foci, while axonal growth is coordinated by a single, dominant spatiotemporal program or by distinct programs that are indistinguishable at the spatial resolution of current diffusion imaging. By explicitly modelling different heterochronous gradients for each neurobiological process (e.g., cell birth, axon growth, myelination) we can test whether different principles really exist.

A major strength of the study is replication across independent cohorts, parcellations, and tractography algorithms, thus increasing our confidence that the heterochronous mechanism is capturing a fundamental organizational principle rather than specific artefacts. Despite the overall gains in capturing network topography (e.g., the location of hubs), the model still falls short in capturing the exact connections we observe in real brains, especially long-range ones. Given the repeated failures in simulating long-range connections specifically (16), it is possible that the locations of these connections are simply not established based upon the whole-brain network principles implemented here. Instead it is possible that the locations of these crucial long-range connections are determined by the need to achieve specific idosyncractic functional goals within our evolutionary past (37). For example, even if the prefrontal-occipital pathway is metabolically expensive, it is essential for top-down modulation of vision, and therefore it might be retained even if it violates the usual principles of wiring economy.

Alternatively, the poor fit of the model to long-distance connections might be due to imprecise modelling or data. On one hand, it is possible that these modelling approaches are still incomplete, and introducing new developmental constraints might enhance our ability to map long-range connections (38). For example, at earlier developmental stages brains are smaller and distances between regions are therefore smaller as well, which may affect the metabolic cost of connections (39) and consequently its trade-off with other generative parameters. Future research should implement these additional biophysical constraints in GNMs. On the other hand, most DTI connectomes underweight the probability of long-distance connections due to well-known issues with crossing fibres and higher false-negative rate for long, tenuous pathways (40, 41). Hence, poorer fit for long connections might reflect limitations in the “gold standard” empirical data rather than shortcomings of the generative model. A possibility for future work would be to test these models in other species such as macaques, where different methods can be applied (e.g. DTI vs tract tracing) and directly compared, while being mindful that these developmental patterns strongly vary across species.

Finally, the heterochronous formulation opens the door to modelling individualised developmental trajectories. This is crucial because network formation processes heavily influence both the structural layout of neural circuits and their functional capabilities, impacting cognitive development as well as psychological wellbeing (42–44). Prior GNMs have been used to probe group differences (10, 14) and individual differences (15), and parameters related to homophily have sometimes been given biological interpretations. However, other terms – most notably the distance penalty – are harder to map onto specific neurodevelopmental mechanisms. In contrast, the parameters we introduce, such as the strength of the heterochronous gradient and its timing, allow for mapping individual differences in best-fitting parameters to clear biological and developmental processes. This opens to the possibility to estimate individualised *changes* in brain organization. Future research should explore how different trajectories in network formation may be linked to diverse network organizations and even variability in cognitive functioning (45, 46).

## Materials and Methods

### Datasets

The Cambridge Centre for Ageing and Neuroscience (CamCAN) study tracks neuro-cognitive changes across the lifespan (47). Conducted at the MRC Cognition and Brain Sciences Unit, University of Cambridge, it focuses on the neural underpinnings of healthy ageing (see Shafto et al., 2014 for full sample and imaging details). All diffusion-MRI gradient directions (i.e., “b-tables”) were validated with an automated quality-control routine to ensure accuracy (48).

The Human Connectome Project young adult (HCPya) dataset was acquired across several sites by the WU-Minn consortium to characterise brain connectivity in a large sample of healthy adults (see Van Essen et al., 2013 for further sample and imaging information). A group-average template was built from 930 participants, and diffusion data were reconstructed in MNI space with q-space diffeomorphic reconstruction (50) to obtain spin-distribution functions (51).

To isolate heterochronicity from broader age-related effects, we confined our analyses to participants aged 26-35 years, thereby avoiding lifespan developmental patterns that could add noise. For this reason, in the present analyses we retained only participants aged 26-35 years, resulting in 78 CamCAN and 829 HCPya individuals. Finally, we also analyzed a developmental sample (CALM) to test the generalizability of the findings; details of acquisition and preprocessing are reported in the Supporting Text.

### Data Preprocessing

The data was accessed in a reconstructed format for DSI Studio’s Fiber Data Hub (52). Each participant’s structural connectome was then traced with *deterministic* tractography based on generalized q-sampling imaging (GQI) in DSI Studio (51), using the *qsiprep* dsi_studio_gqi workflow (53). Tractography was seeded in all voxels of the AAL-116 atlas (54); subcortical parcels 91–116 were then excluded, yielding the AAL-90 cortical parcellation (78 cortical areas and 12 subcortical/limbic areas). We selected this atlas because its well-validated 90-region resolution consistently recovers canonical connectome features (e.g., small-worldness, rich-club hubs, see 31–33). To control for atlas differences, we applied the same data preprocessing also using the Schaefer100 atlas (58).

Tracking parameters were identical for every participant: 5 million streamlines, seeded at random; step size = 1 mm; turning angle threshold = 35°; minimum / maximum fibre length = 30 mm / 250 mm. Connectivity was quantified with a count-end definition: a streamline contributed to the connection matrix only if it terminated in both of the paired regions. The resulting edge weight (our variable of interest) was the streamline count between each pair of parcels.

Once individual connectomes were obtained, we built a consensus connectome using the *struct_consensus* function from the *netneurotools* package in Python, which relies on the distance-dependent algorithm as implemented by Betzel and colleagues (32). Once a consensus network was created for each dataset, we thresholded it to 10% density based on the strongest connections (15). Finally, the networks where binarised. The resulting consensus networks were in the form of single-component adjacency matrices. The same connectomes were also reconstructed with the Schaefer100 parcellation and, for CALM, with probabilistic tractography; these variants and their thresholding are described in the Supporting Text.

### Computational modelling

Models were generated following equation (4) in the Results section. *K, D* and *H* represent the matrices for homophily (i.e., matching index), distance, and heterochronicity, respectively. Homophily was computed as the matching index of the adjacency matrix, which is a measure that quantifies the similarity of input and/or output connections of two nodes excluding their mutual connections (10, 59, 60); Euclidean distance was computed simply by calculating the straight-line distance between the spatial coordinates of node pairs in the anatomical space; Heterochronicity was computed using a time-varying cumulative Gaussian function. More specifically, the likelihood of a certain node *i* to establish any connection at the time *t* was given by a Gaussian distribution:

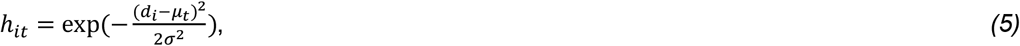

where *d*_*i*_ is the Euclidean distance of node *i* from the origin point, σ is the standard deviation of the Gaussian (which was kept at a fixed value, σ = 15) and µ_*t*_ changes across time as follows:

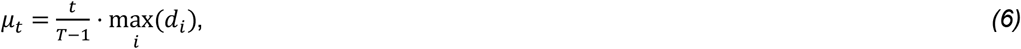

thus placing the centre of the Gaussian µ_*t*_ at equally spaced positions along the distance axis from the reference point out to the farthest node. A temporal scaling parameter τ further regulated the timing of this developmental sweep: τ = 1 yields the linear progression of equation (6); τ < 1 accelerates the wave so it reaches the periphery early and stays fully unfolded thereafter; τ > 1 slows it so the Gaussian centre never reaches the farthest node within T steps. For the main analyses τ was held fixed at 1; in the individual-differences analysis (Fig. S2) it was allowed to vary across participants.

Once a node became “active” (*a*) and started making connections, it remained active until the end of the heterochronous process, such that:

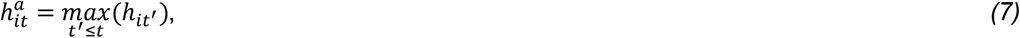

where *t*^′^ indexes over all previous time steps up to and including *t*. Once the likelihood of connecting was defined for each node, an undirected matrix could be generated, thus specifying the likelihood of any two nodes *i* and *j* of connecting:

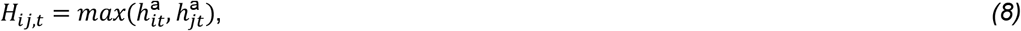

A simple representation of this process can be found in Fig. 1D. When dual origins were introduced, the process was now characterised by two independent origin points, which were then summed and normalised to obtain a single matrix *H*_*ij,t*_. Hence, dual origin models had the same temporal process as single orgin models, rather than differently paced waves of heterochronous development.

### Model fitting

The models were fitted to the consensus networks to minimize the topological and topographical energies. In the standard energy function, only topology is taken into account. More specifically, the difference in the cumulative distributions of four network metrics between real and synthetic networks is computed in terms of Kolmogormov-Smirnoff (KS) similarity, such that the energy of the synthetic model is computed following eq. 1. In addition, we also considered a topological energy. To compute this energy, we calculated the correlation *r* between the synthetic and actual nodal metrics (i.e., nodal degree, betweenness centrality, clustering coefficient), and took the smallest value as the energy (eq. 2). The resulting total energy was a sum of topological and topographical energies (eq. 3).

Given that we aimed to capture the *approximate* location of the network properties, we performed a spatial smoothing before computing the topographical energy. For a node-level network property X_i_, the unnormalised spatial smoothing weights 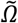 are given by:

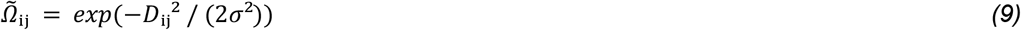

For each node *i*, a larger weight is assigned to those nodes *j* which are closer to *i* in space (i.e., have a lower value of *D*_*ij*_). The parameter *σ* controls the spatial scale of the smoothing. These weights do not sum to 1, so we normalise by performing:

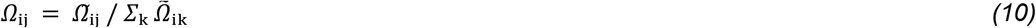

The node-level property is then smoothed using these weights to obtain σ_j_ Ω_ij_ X_j_. The correlation between these smoothed quantities across the real and synthetic networks can then be computed.

When fitting the model, γ, η, and λ (which regulate the impact of homophily *K*, Euclidean distance *D*, and heterochronicity *H*, respectively on the probability of making a connection) were free parameters. More specifically, each of them was free to vary as follows: γ = [0, 0.8], η = [-3.5, 0], and λ = [0, 15]. The ranges of γ and η were based on previous literature (15) while the range of λ was kept broader because no prior knowledge is available.

In addition, the coordinates of the origin point of the heterochronous gradient were also treated as free parameters. In the first analysis, the range of values was defined following the medial plane. First, we found the minimum and maximum *y* and *z* values across all brain coordinates to create a 3D box that fully contains the brain. Next, we sample 9 points in each direction (i.e., 9 along the y axis and 9 along the z axis), evenly spacing sample points across each axis within the bounding box, forming a 2D grid of points. Finally, we checked which of those grid points actually fall inside the brain’s shape (not just the box) using a *convex hull* — a geometric wrapper that approximates the outer surface of the brain. The points that fell inside the brain were used as potential origins for heterochronous development. In the subsequent analyses (Fig. 4), we used the nodes of the AAL atlas as origin points.

A grid-search was performed over the multidimensional parameters space. For parameters γ, η, and λ, 10 equidistant values were sampled. The medial sweep approach returned 42 valid origin points. This resulted to a total of 42000 parameters combinations. Given that network formation is a stochastic process, we run each combination of parameters 10 times. All network metrics reported in the paper are averaged across runs to reduce noise in the estimates.

The combination of parameters that lead to the lowest energy was similar for the camCAN and the HCPya datasets. For camCAN, the lowest Energy_total_ was 0.296 (with Energy_tolopogy_′ = 0.199 and Energy_topography_′ = 0.395). The lowest energy was obtained with parameters values η = -1.17, γ = 0.23, and λ = 5.0, and with origin point coordinates y = -66.97 and z = 44.03. For HCPya, the lowest Energy_total_ was 0.315 (with Energy_tolopogy_′ = 0.446 and Energy_topography_′ = 0.187). The lowest energy was obtained with parameters values η = -1.56, γ = 0.23, and λ = 5.0, and with origin point coordinates y = -66.97 and z = 31.01.

### Analysis

All statistical analyses were carried out in R (version 4.2.2), using the *lm* function from the *stats* package and the *gam* function from the *mgcv* package. Results for HCPya and CamCAN with the AAL parcellation are reported in the main text; all remaining reconstructions (Schaefer100, CALM, and probabilistic tractography) are reported in Fig. S4-S5.

## Acknowledgments

All data provided on DSI Studio were analysed using the US National Science Foundation, ACCESS program, resource allocation (TG-CIS200026) at Extreme Science and Engineering Discovery Environment (XSEDE) resources (Towns, J. et al. Computing in science & engineering 16, 62-74 2014) by Fang-Cheng Yeh at the University of Pittsburgh.

## Author contributions

F.P.: Conceptualization, Methodology, Formal analysis, Data Curation, Writing – Original draft, Visualisation, Funding acquisition; S.O.: Conceptualization, Writing – Review & Editing; A.M.: Data Curation, Writing – Review & Editing; E.T.B.: Conceptualization, Writing – Review & Editing; P.E.V.: Conceptualization, Writing – Review & Editing; D.A.: Conceptualization, Methodology, Resources, Writing – Review & Editing, Supervision, Funding acquisition.

## Data and materials availability

All data needed to evaluate the conclusions in the paper are present in the paper and/or the Supporting Information. Data, code, and materials are available on OSF: https://osf.io/2yuh7.

