## Supplementary Information for "Heterochronicity shapes brain network formation"

PNAS

*Francesco Poli

**This PDF file includes:**

Supporting text

Figures S1 to S6

### Supporting text

To test the generalizability of our findings, we extended our analyses to a different parcellation, a different dataset, and, within the latter, also a different tractography method (probabilistic). In addition to the original AAL90 (90 regions) analyses of the adult HCPya and CamCAN cohorts, we re-derived group-representative connectomes using the 100-region Schaefer atlas (Schaefer100), and we introduced a developmental sample, the Centre for Attention, Learning and Memory (CALM) cohort (age range 6.1-18.8 years, $M = 10.3$, $SD = 2.3$). Data acquisition and pre-processing steps for the CALM dataset can be found in Mareva et al. (2022).

For CALM we reconstructed connectomes with both deterministic ($N = 392$) and probabilistic ($N = 387$) tractography, letting us additionally assess robustness to the tractography approach. Deterministic networks were thresholded to a 10% group-consensus edge density, matching the main analyses; probabilistic networks, which support a denser reliable consensus, were thresholded to 20%. In total, this yielded six group-representative connectomes spanning three datasets (HCPya, CamCAN, CALM), two parcellations (AAL90, Schaefer100), and two tractography methods: AAL90/deterministic for HCPya and CamCAN; Schaefer100/deterministic for HCPya; and, in CALM, AAL90/deterministic, Schaefer100/deterministic, and Schaefer100/probabilistic. The generative model, energy function, and parameter grid were held identical across all reconstructions, so that results overlay directly.

The same analyses were applied to all six group-representative connectomes; they are reported for HCPya in Fig. S4 and for CALM in Fig. S5.

*Parameters contributions.* To establish that each model term contributes to fit rather than being absorbed by the others, we took each reconstruction's best-fitting heterochronous model and re-ran it with a single parameter set to zero while holding the other two at their optimum: $\eta = 0$ removes the distance term, $\gamma = 0$ removes the matching-index term, and $\lambda = 0$ removes the heterochronicity term, recovering the classical generative model. Each configuration was simulated 10 times, and each ablated model was compared with the full model by Welch's t-test on total energy.

*Connection-length distributions.* For each of 30 simulation runs per model we extracted the Euclidean length of every generated edge and compared the resulting distribution against that of the empirical consensus network using the KS statistic. Because synthetic and empirical networks are matched in edge count, this compares distributional shape only. KS distances were then compared between the heterochronous and classical models by Welch's t-test across runs.

*Origin position.* The heterochronous wave was seeded at each of 42 candidate origins spanning the brain. Holding $\eta$, $\gamma$ and $\lambda$ at each reconstruction's best-fitting values, we modelled total energy as a smooth function of the origin's y-coordinate (anterior–posterior position) with a generalised additive model (thin-plate spline, $k = 4$, REML), marginalising over the z-coordinate; with 10 runs per origin this gives $n = 420$ observations per reconstruction. We report the effective degrees of freedom, F, p and deviance explained of the smooth term, together with the position of the fitted optimum.

*Edge placement by distance.* To ask where each model places its edges correctly, we recorded for every node pair the probability with which the model placed an edge there, spatially smoothed with the same kernel used for the topographical energy. Placement probability was then modelled as a function of connection length with a generalised additive model containing separate smooths ($k = 6$) for node pairs that are connected and unconnected in the empirical consensus network. A model that places edges where the real network has them produces a higher curve for present than for absent pairs; a model performing at chance produces coincident curves. The difference between the two smooths was evaluated with a simultaneous 95% confidence band, which controls error across the whole distance range, and the interval over which this band excludes zero is drawn in bold.

### Figures

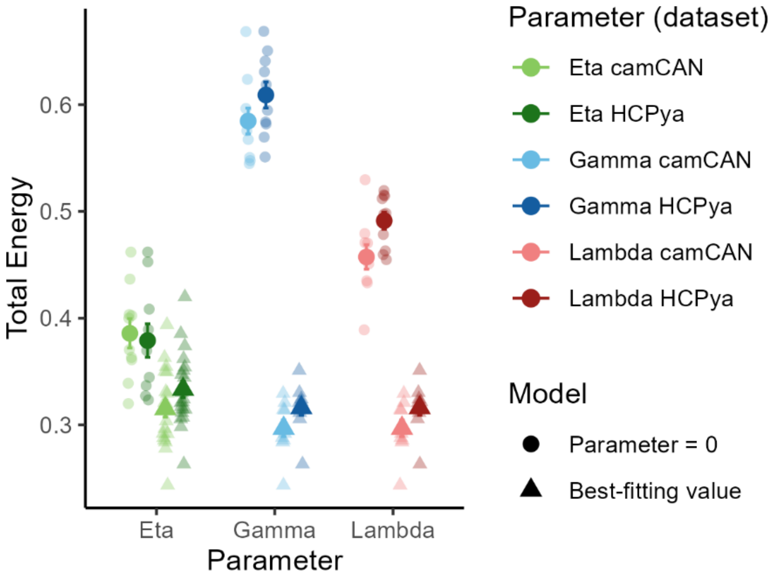


**Fig. S1.** Total energy (lower = better fit) of generative network models fitted to camCAN (light shades) and HCP-YA (dark shades) connectomes, shown separately for the three model parameters: η, the distance term (green); γ, the matching-index term (blue); and λ, the heterochronicity term (red). In each case the best-fitting model (triangles) is compared against the same model with that one parameter set to zero (circles), i.e. with the corresponding term removed while the other two remain at their optimum. Small semi-transparent symbols show the 10 independent stochastic simulations (i.e. runs); large symbols and error bars show the mean ± SEM. Removing the matching term produces the largest loss of fit, followed by heterochronicity and then distance.

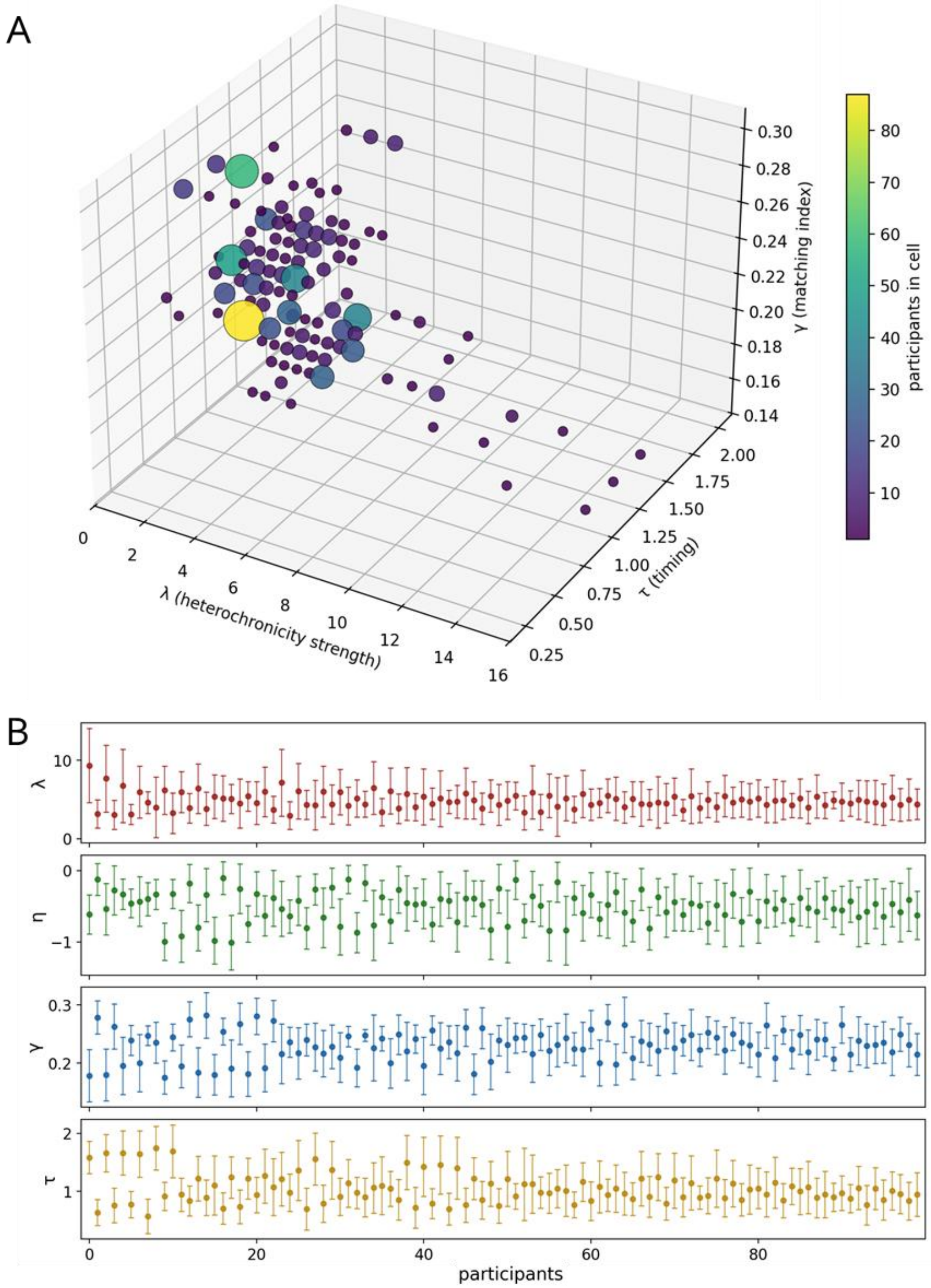


**Fig. S2.** Individual variability in best-fitting GNM parameters (AAL90, HCP-YA), with N = 20 stochastic runs per combination, with 16 × 11 × 8 × 8 grid of (λ, τ, η, γ) combinations, scored against each participant via the topology + topography energy. **A.** 3-D heatmap of each participant's lowest-energy (λ, τ, γ) combination (η profiled out by using each participant's own best-fitting η); marker colour and size denote the number of participants whose best combination falls in that cell. **B.** 100 participants selected to display individual differences, shown as per-participant mean ± SD for each parameter and ordered along the x-axis so that adjacent participants are maximally different.

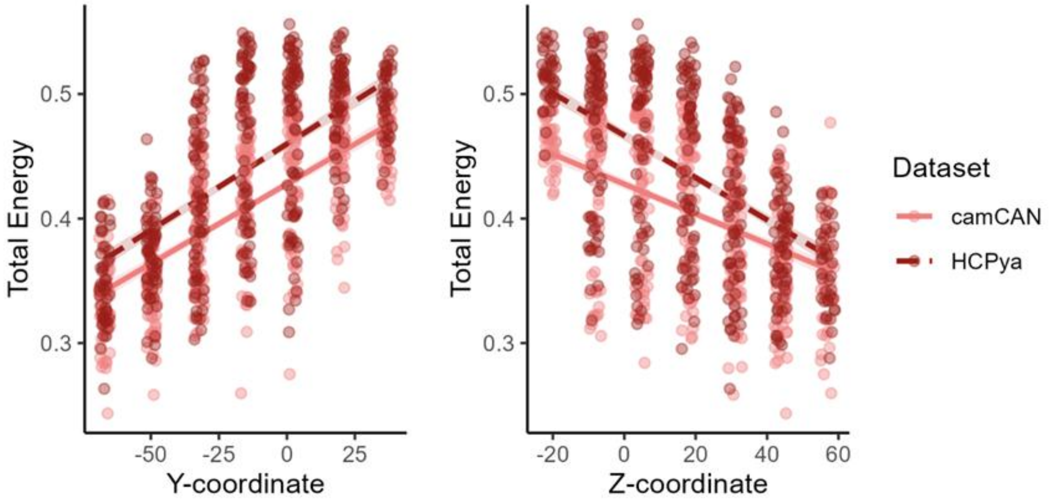


**Fig. S3.** Scatterplots depicting total energy values as a function of Y (left) and Z (right) coordinates. Lower energy is obtained with lower Y coordinates and higher Z coordinates.

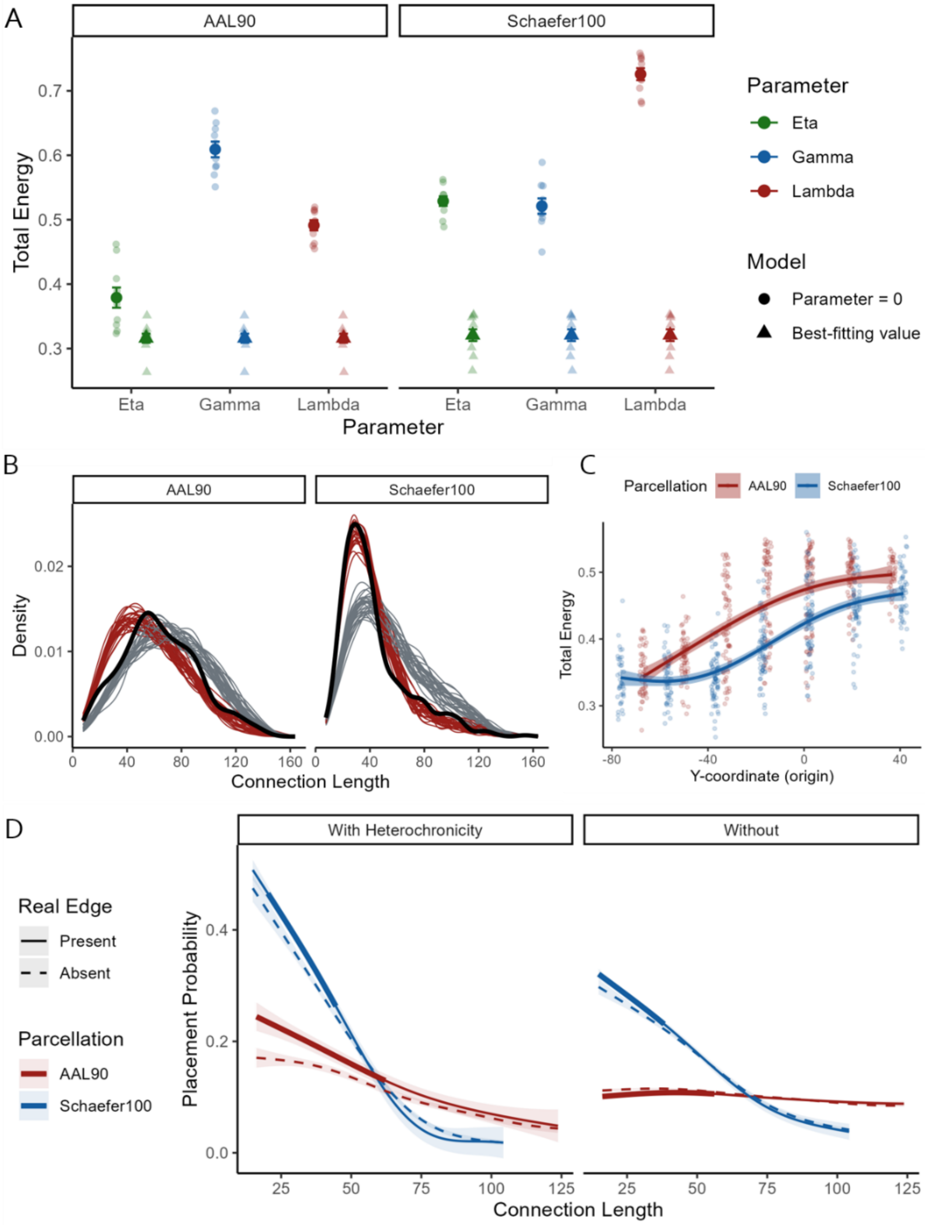


**Fig. S4.** All panels compare the AAL90 (90-region) and Schaefer100 (100-region) parcellations of the adult HCPya connectome, with deterministic tractography throughout. **A.** Total energy (lower = better fit) of generative network models under the two parcellations, AAL90 (left) and Schaefer100 (right), shown separately for the three model parameters: η, the distance term; γ, the matching-index term; and λ, the heterochronicity term. In each case the best-fitting model is compared against the same model with that one parameter set to zero, i.e. with the corresponding term removed while the other two remain at their optimum. Small semi-transparent symbols show the 10 independent stochastic simulations (i.e. runs); large symbols and error bars show the mean ± SEM. Removing any of the three terms substantially degrades fit in both parcellations (all $p \leq .003$, Welch's t-test). **B.** Distributions of connection lengths generated by the heterochronous model (red) and the classical model (grey) across 30 individual simulation runs, plotted against the empirical consensus network (black), for AAL90 (left) and Schaefer100 (right). In Schaefer100 the heterochronous model reproduces the empirical distribution markedly better than the classical model (Kolmogorov–Smirnov distance to the consensus distribution, mean ± SD across runs: 0.054 ± 0.013 vs 0.255 ± 0.030; Welch $t(39.4) = 33.56$, $p < 0.0001$, Hedges' $g = 8.55$), with the classical model over-producing long connections (median +13.7 mm relative to the consensus, vs +1.5 mm for the heterochronous model). In AAL90 the two models fit the empirical distribution equally well overall (0.135 ± 0.024 vs 0.130 ± 0.024; $t(58.0) = -0.78$, $p = .44$), but deviate in opposite directions: the heterochronous model generates connections that are somewhat too short (median −6.9 mm) whereas the classical model generates connections that are too long (+8.7 mm). **C.** Total energy as a function of the y-coordinate (anterior–posterior position) of the heterochronous wave's origin. Each point is one simulation run at one of the 42 candidate origins, with $\eta$, $\gamma$ and $\lambda$ held at that parcellation's best-fitting values ($n = 420$ runs per parcellation); the curve is a GAM smooth (thin-plate spline, $k = 4$) with its 95% confidence band, and is therefore marginal over the z-coordinate. Fit improves markedly as the origin moves posteriorly in both parcellations (AAL90: edf = 2.61, $F = 135.5$, $p < 0.0001$, 48.4% deviance explained; Schaefer100: edf = 2.92, $F = 183.2$, $p < 0.0001$, 56.9%), with the fitted optimum at $y \approx -67$ mm and −59 mm respectively. **D.** Probability of placing an edge as a function of connection length, shown separately for node pairs that are present (solid) and absent (dashed) in the empirical consensus network, for the heterochronous (left) and classical (right) models. The heterochronous model places edges preferentially where the real connectome has them at short range in both parcellations (AAL90: present > absent from 16 to 62 mm, maximum difference +0.074 at 16 mm; Schaefer100: 20–45 mm, maximum +0.033), converging to chance for longer connections. The classical model shows no comparable short-range accuracy.

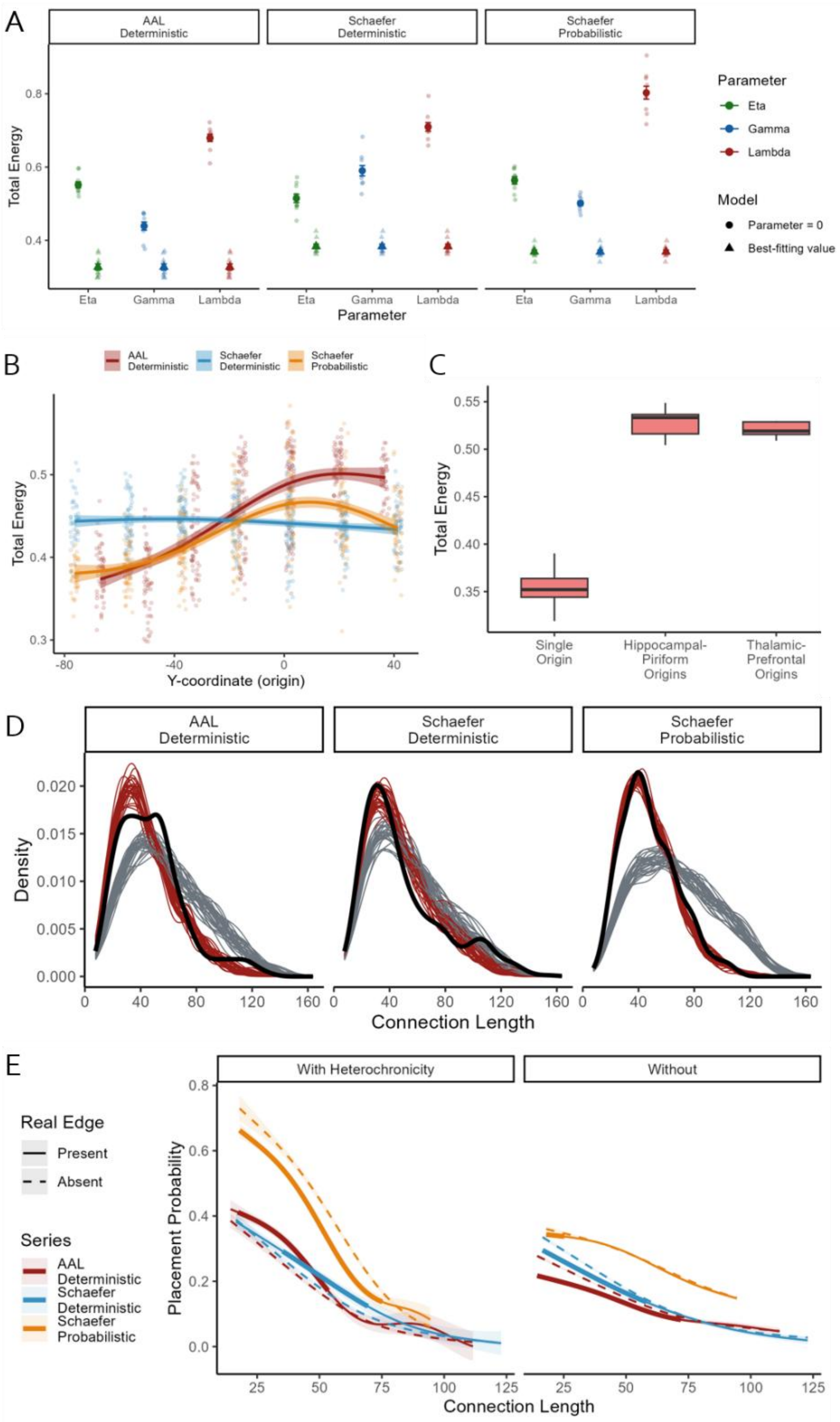


**Fig. S5.** Generalizability of the heterochronous model for the CALM dataset across AAL90 with deterministic tractography, Schaefer100 with deterministic tractography, and Schaefer100 with probabilistic tractography. **A.** Total energy (lower = better fit) of generative network models under the three parcellation × tractography combinations: AAL90 deterministic (left), Schaefer100 deterministic (middle) and Schaefer100 probabilistic (right), shown separately for the three model parameters η, γ and λ. In each case the best-fitting model (triangles) is compared against the same model with that one parameter set to zero (circles), i.e. with the corresponding term removed while the other two remain at their optimum. Small semi-transparent symbols show the 10 independent stochastic simulations (i.e. runs); large symbols and error bars show the mean ± SEM. Removing any of the three terms substantially degrades fit in all three reconstructions (all $p \leq .001$, Welch's t-test). **B.** Total energy as a function of the y-coordinate (anterior–posterior position) of the heterochronous wave's origin. Each point is one simulation run at one of the 42 candidate origins, with η, γ and λ held at that series' best-fitting values (n = 420 runs per series); the curve is a GAM smooth (thin-plate spline, k = 4) with its 95% confidence band, marginal over the z-coordinate. A posterior origin yields substantially better fits under AAL-deterministic (edf = 2.87, F = 149.2, $p < 0.0001$, 51.7% deviance explained; fitted optimum at y ≈ −67 mm, with energy rising by 0.13 towards anterior origins) and Schaefer-probabilistic (edf = 2.95, F = 85.7, $p < 0.0001$, 38.6%; optimum at y ≈ −76 mm, rise of 0.09), whereas under Schaefer-deterministic the anterior–posterior position of the origin has essentially no effect (edf = 2.05, F = 4.0, p = 0.021, 2.4% deviance explained; fitted energy varies by only 0.013 across the full range). **C.** Total energy of the heterochronous generative model under three origin configurations: a single bilateral caudal origin (the configuration identified by the origin sweep), a bilateral hippocampal–piriform dual origin, and a bilateral thalamic–prefrontal dual origin. η and γ were fixed at CALM's best-fitting values (η = −3.11, γ = 0.11) and λ was swept from 0 to 15, with each configuration evaluated at its own best-fitting λ over 10 independent stochastic simulations; boxes show the median and interquartile range, whiskers extend to 1.5 × IQR. Both theory-driven dual origins fit substantially worse than the single caudal origin (hippocampal–piriform 0.528 ± 0.015 and thalamic–prefrontal 0.526 ± 0.016 vs 0.352 ± 0.021, mean ± SD; linear model with the single origin as reference: β = 0.176, SE = 0.008, t(27) = 22.30, $p < 0.0001$ and β = 0.174, SE = 0.008, t(27) = 22.02, $p < 0.0001$; model $R^2 = 0.96$). This analysis is presented for the AAL90 parcellation only: the Schaefer100 parcellation is cortical-only and contains no hippocampal, piriform or thalamic parcels, so the subcortical origin points required by the two dual-origin theories cannot be placed in it. **D.** Distributions of connection lengths generated by the heterochronous model (red) and the classical model (grey) across 30 individual simulation runs, plotted against the empirical consensus network (black), for the three reconstructions. The heterochronous models reproduce the empirical distribution better than the classical models in every case (Kolmogorov–Smirnov distance to the consensus distribution, mean ± SD across runs: AAL-deterministic 0.106 ± 0.020 vs 0.202 ± 0.028, Welch t(53.1) = 15.25, $p < 0.0001$; Schaefer-deterministic 0.079 ± 0.010 vs 0.158 ± 0.026, t(37.6) = 15.31, $p < 0.0001$; Schaefer-probabilistic 0.040 ± 0.010 vs 0.289 ± 0.028, t(36.6) = 45.95, $p < 0.0001$; all Hedges' g ≥ 3.9), the classical model systematically over-producing long connections. **E.** Probability of placing an edge as a function of connection length, shown separately for node pairs that are present (solid) and absent (dashed) in the empirical consensus network, for the heterochronous (left) and classical (right) models. Under AAL-deterministic and Schaefer-deterministic the heterochronous model places edges preferentially where the real connectome has them at short to intermediate range (present > absent over 17–53 mm, maximum difference +0.077 at 34 mm; and over 35–69 mm, maximum +0.037 at 55 mm), converging to chance at longer distances, whereas the classical model shows the opposite pattern in both (present < absent over 14–72 mm and 17–58 mm, maximum magnitudes 0.061 and 0.040). Under Schaefer-probabilistic neither model shows short-range accuracy, both falling below their absent counterparts (heterochronous 18–75 mm, maximum magnitude 0.086; classical 18–25 mm, maximum 0.017). However, for probabilistic tractography, the location and strength of the hubs is unreliable (see Fig. S6). This suggests that failure in this case is due to limitations in the data rather than the model.

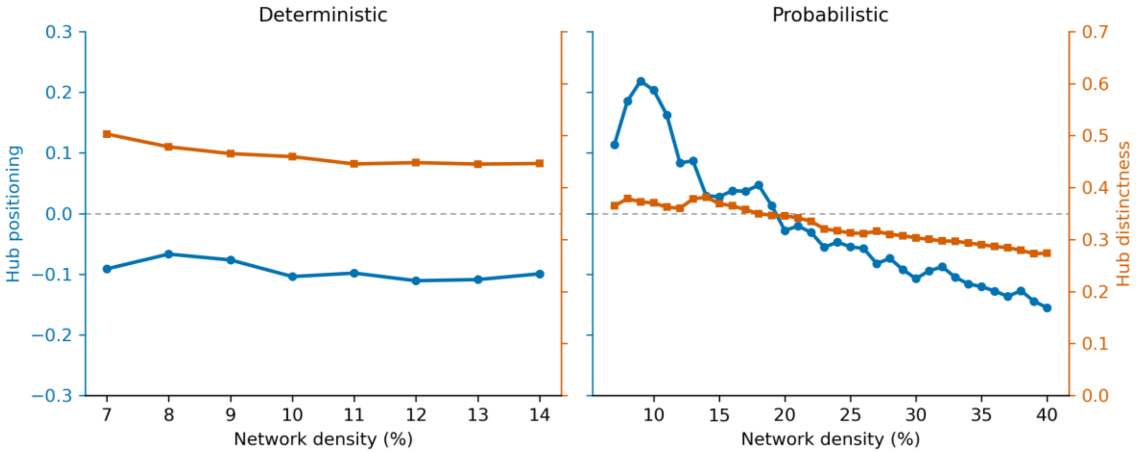


**Fig. S6.** Hub organisation of the CALM Schaefer100 empirical connectome as a function of network density, for deterministic (left; 7-14%) and probabilistic (right; 7-40%) tractography. Each network was obtained by retaining the strongest edges of the group consensus down to the density on the x-axis. Two properties of the resulting degree distribution are shown. Hub positioning (blue, left axis) is the Spearman correlation between a node's degree and its y-coordinate (anterior-posterior position). Hub distinctness (orange, right axis) is the coefficient of variation of the degree distribution: higher values indicate a more heavy-tailed distribution with a small number of clearly differentiated high-degree hubs, lower values a flatter distribution in which hubs are weakly differentiated. Under deterministic tractography, both properties are stable across the density range: hubs remain caudally positioned (blue ≈ −0.10) and distinct (orange ≈ 0.45-0.50) at every density, including the 10% used for modelling. Under probabilistic tractography, neither property is stable.
